# POPSHIFT: A THERMODYNAMICALLY SOUND APPROACH TO ESTIMATE BINDING FREE ENERGIES BY ACCOUNTING FOR LIGAND-INDUCED POPULATION SHIFTS FROM A LIGAND-FREE MSM

**DOI:** 10.1101/2023.07.14.549110

**Authors:** Louis G. Smith, Borna Novak, Meghan Osato, David L. Mobley, Gregory R. Bowman

## Abstract

Obtaining accurate binding free energies from *in silico* screens has been a longstanding goal for the computational chemistry community. However, accuracy and computational cost are at odds with one another, limiting the utility of methods that perform this type of calculation. Many methods achieve massive scale by explicitly or implicitly assuming that the target protein adopts a single structure, or undergoes limited fluctuations around that structure, to minimize computational cost. Others simulate each protein-ligand complex of interest, accepting lower throughput in exchange for better predictions of binding affinities. Here, we present the PopShift framework for accounting for the ensemble of structures a protein adopts and their relative probabilities. Protein degrees of freedom are enumerated once, and then arbitrarily many molecules can be screened against this ensemble. Specifically, we use Markov state models (MSMs) as a compressed representation of a protein’s thermodynamic ensemble. We start with a ligand-free MSM and then calculate how addition of a ligand shifts the populations of each protein conformational state based on the strength of the interaction between that protein conformation and the ligand. In this work we use docking to estimate the affinity between a given protein structure and ligand, but any estimator of binding affinities could be used in the PopShift framework. We test PopShift on the classic benchmark pocket T4 Lysozyme L99A. We find that PopShift is more accurate than common strategies, such as docking to a single structure and traditional ensemble docking—producing results that compare favorably with alchemical binding free energy calculations in terms of RMSE but not correlation—and may have a more favorable computational cost profile in some applications. In addition to predicting binding free energies and ligand poses, PopShift also provides insight into how the probability of different protein structures is shifted upon addition of various concentrations of ligand, providing a platform for predicting affinities and allosteric effects of ligand binding. Therefore, we expect PopShift will be valuable for hit finding and for providing insight into phenomena like allostery.

## I Introduction

Developing strategies to accelerate and simplify hit discovery in drug development is one of the core foci of computational chemistry. Because huge arcs of chemical space must be subtended, methods that scale well per ligand predominate.^1^ Most of these methods are based on docking a set of compounds to a single protein structure as rapidly as possible to maximize the chemical space that can be considered. The scores predicted by these methods correlate so poorly with true binding affinities that they are typically judged by how much the high scoring compounds are enriched for tight binders compared to randomly selected compounds.^2,3^ Of course, a wide range of methods have been developed to make different trade-offs between speed and accuracy. Of these, alchemical free energy calculations are some of the most physically rigorous and should, in principle, be capable of quantitatively accurate predictions.^4,5^ However, routinely achieving quantitative predictions with any method remains difficult.^6^

One striking feature of all these methods is the extent to which they assume proteins adopt a limited set of highly similar structures. Many docking algorithms don’t include any protein conformational heterogeneity. The cross docking problem highlights the limitations this assumption imposes (i.e. docking a library of compounds against a protein structure obtained by removing a ligand from a ligand-bound structure is more predictive than docking against a structure obtained in the absence of ligand).^7^ To address this, some docking algorithms allow limited protein flexibility, such as rotations of side-chains. However, many do not find that incorporating conformational heterogeneity in this way is worth the additional computational cost.^8^ In principle, alchemical free energy calculations should be able to deal with protein conformational heterogeneity as every degree of freedom is allowed to move as dictated by the force field. However, in practice, alchemical free energy simulations are so short that the protein only undergoes limited fluctuations around the starting structure.^9^ Phrased differently, ignoring receptor conformational heterogeneity for the sake of computational performance is one of the key approximations of most digital screening campaigns.

Ensemble docking has emerged as a strategy to address protein conformational heterogeneity but still faces significant limitations.^10^ In ensemble docking, one generates a set of protein structures (often via molecular dynamics simulations) and then docks a library of compounds against each of these structures. Typically, one then ranks the compounds based on their best score against any protein structure, though there are other flavors of ensemble docking. While this ensemble docking approach recognizes there is uncertainty in which protein structure is relevant, it still essentially assumes that a single structure is relevant in the end. It also throws out thermodynamic information from the simulations, instead giving all protein structures equal weight. These methods are still generally incapable of quantitative predictions and suffer from some strange pathologies. For example, it has been reported that ensemble docking against short simulations outperforms docking to a single structure but that adding more simulation data often hurts performance rather than helping.^10–13^ Other efforts to include conformational heterogeneity into docking have included the existence of multiple conformations using some assessment of their relative abundance, but have done so in an ad-hoc fashion.^14,15^

Here, we propose a reweighting approach called PopShift that uses Markov state models (MSMs) of a ligand-free protein to account for the populations of different protein structures and how they are shifted upon binding to a ligand. MSMs can be viewed as a compressed representation of the system’s thermodynamic ensemble.^16^ Thus, MSMs representing the ligand-free protein ensemble contain all the receptor information needed to estimate ligand binding.^17^ In order to weight the contribution of state populations in the apo context versus how tightly they bind ligand, we estimate binding to representative conformations from each state, obtaining a per-state binding free energy by averaging them. We then use a binding polynomial approach to track which states actually contribute to the macroscopic affinity for the ligand that might be observed in an experiment like ITC or similar.^18^ Equivalently, the presence of the ligand can be viewed as a perturbation to the state in the sense of the Zwanzig formula.^19^ Thus, instead of taking the best score against any structure, as in traditional ensemble docking, we take a correctly weighted average over all structural states. This is made tractable by the MSM, since it means we only need one or, to be more confident, a handful of binding estimates per MSM state. A related idea that treats affinity per conformation using the same math, but does not leverage an MSM to index conformational heterogeneity, is the Implicit Ligand Theory.^20,21^

By capturing how ligands shift the relative probabilities of different protein conformations, PopShift also provides an opportunity to understand how ligands remodel their binding sites or even allosterically impact distant sites. Population-shift in response to ligand binding is, by definition, allostery. If the ligand-free MSM is a compressed representation of the perturbed model’s ensemble, then the reweighted MSM is a compressed representation of the receptor’s liganded ensemble. Thus, observables of interest can be estimated with the reweighted state probabilities to understand allosteric mechanism. Because the expression for reweighted state probabilities includes ligand concentration, the impact on these averages can also be used to compute an EC50.

To test PopShift, we compare its performance to several other candidates on a simple-yet-subtle benchmark for protein-ligand binding, T4 Lysozyme L99A. From a set of apo simulations we build an MSM. We then sample conformations from each MSM bin and dock to them, using the customary organic fragments from Morton et al. [22]. We use the docking score as a heuristic for the free energy of binding to a particular conformation. We recognize that docking has severe limitations, especially for ligands with rotatable bonds. However, docking provides a simple and highly relevant starting point given its widespread use in drug discovery, and the fragments we consider here are not subject to the known issues with rotatable bonds. In the future, it will be interesting to try alternatives to docking in the PopShift framework. In PopShift, the free energies of binding are then incorporated into an affinity estimate and reweighted state probabilities using PopShift. We compare these strategies to best-score docking, docking to holo crystal structures with the ligand removed, and to absolute binding free energy calculations performed in the customary style with docked and hand-adjusted starting poses. We also explore how the conformational preferences of the protein are altered by the addition of ligand.

## II Results and Discussion

### II.A PopShift performs well compared to alternative in silico estimators of binding free energies

We reasoned that modern simulations are sufficiently predictive that both structures from these simulations and their populations can inform a successful hit finding strategy. In particular, MSMs provide a powerful and quantitatively predictive map of a protein’s conformational ensemble—and therefore approximate its partition function. Thus, we hypothesized that using the populations from an MSM in a binding polynomial approach–with the MSM as an approximate partition function–would allow correct incorporation of docking scores from across this sample.

To test this hypothesis, we collected three replica simulation datasets of L99A, made MSMs from them, and estimated binding affinities using PopShift and other popular alternatives. Each replica consisted of 10 simulations, 5×4 μs and 5×8 μs, started from PDB 187L with the ligand (p-xylene) removed. One MSM was made for each replica dataset using TICA on pocket residue backbone and sidechain torsions, and VAMP-2 to validate the number of clusters for k-means as has been done in Meller et al. [23]. We docked ligands from the classic Morton et al. [22] set against structures from each MSM state using the SMINA docking algorithm.^24^ Macroscopic binding affinities were estimated using the PopShift framework, see methods (Section IV). For comparison to extant approaches, we used the conventional ensemble docking approach of taking the best score across a set of samples, and of docking to a crystal structure with ligand removed. We also performed absolute binding free energy simulations using a vanishing ligand transformation from initial hand-selected ligand poses.

We find that PopShift performs well compared to alternative docking approaches and even showed some advantages compared to alchemical free energy calculations (Fig. 2). Docking each ligand to a single holo structure, (the n-butylbenzene structure, PDB 186L) then taking the minimized score as an affinity estimate gives a poor correlation to experimentally measured binding affinities and poor accuracy, as measured by the root mean squared error (RMSE) from experimental results. PopShift also outperforms simply taking the best score, a traditional ensemble docking approach, most notably for ranking. The best-score approach systematically predicts affinities that are too favorable. Although using docking to a ligand-removed crystal structure’s scores as affinity estimates exhibited slightly better correlation with experiment on this dataset than the best-score approach, these scores emanating from a lone structure exhibited a similar pattern of overly favorable affinity estimates. Absolute binding free energy estimates produced the strongest correlation and ranking results, but struggled with accuracy. This is likely related to initial poses and receptor conformations failing to relax fully in the course of the windowed simulations. This interpretation is complicated by our use of the docking energy function to make affinity estimates, which is very different from the forcefield we used to obtain the MSM we dock to. Because docking to many samples from an MSM allows us to estimate affinity to many receptor conformations, it sidesteps the issue of having to chose a ‘most relevant’ one to start from. This is especially important if there may be multiple thermodynamically relevant poses for the ligand.^25^

**Figure 1.**
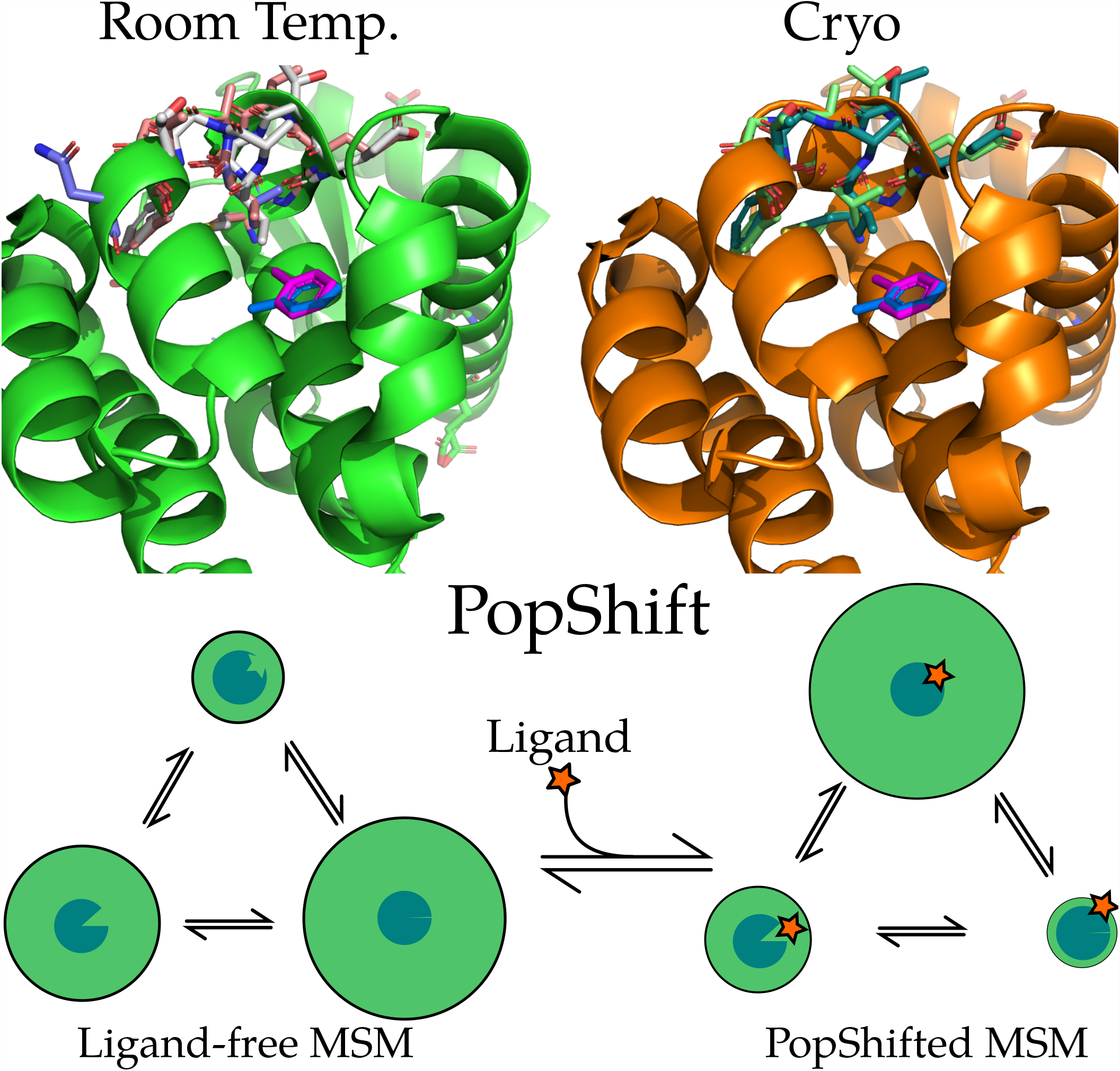
Examples of conformational heterogeneity in T4 lysozyme and a schematic of how PopShift accounts for this heterogeneity. These renders show the multiple conformations even the L99A pocket bound to toluene is capable of accessing under crystallographic study. The top section shows the room temperature structure (PDB 7L39) and shows a cryogenic structure from the same study (PDB 7L3A). All residues with alternative locations in the F-helix, and also toluene, are shown in sticks. Extensive alternative locations are present in both, even though this protein is reckoned to be rigid and to bind simple, largely rigid, fragments. Note the two alternative locations for the ligand are nearly identical at both temperatures. Nearly every residue in the critical F-helix shows heterogeneity, centered on valine 111, which extends down toward the toluene. The lower panel shows a schematic of the PopShift method, showing MSM populations from a ligand free ensemble being biased by varying degrees of ligand affinity to those states, to approximate the ligand-bound ensemble. The sea-blue pac-man represents the protein, with three states in equilibrium and green circle sizes indicating abundance, and the shape cut out of the pac-man representing varying degrees of pocket accessibility to the ligand, which is schematically represented by a star.

**Figure 2.**
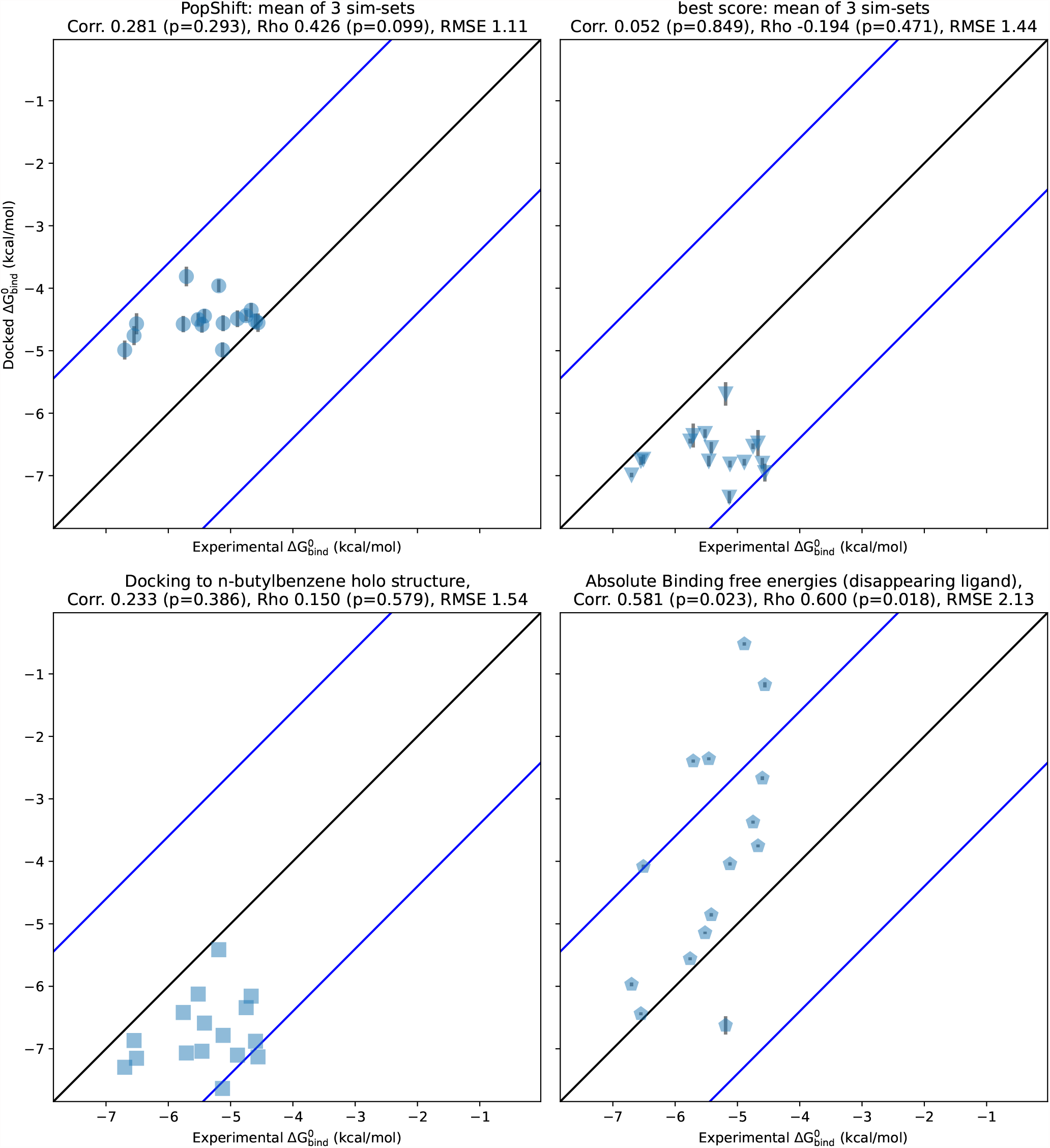
PopShift compares well to alternative predictors, such as docking to a single crystal structure, traditional ‘best score’ ensemble docking, and alchemical free energy calculations. The *x* values are the experimental binding free energies for 17 ligands as measured by ITC in Morton et al. [22]. The y values are the binding free energy estimated by each in silico method. Correlation is the Pearson’s correlation coefficient, rho is the Spearman’s ranking coefficient, and RMSE is the root-mean-squared error in kcal/mol. The error bars on the top two panels are the SEM across the three largest

Based on reports of poorer performance of best score aggregation on longer simulations, we were curious to know how sensitive our method is to dataset size. We reasoned that larger datasets will typically include some incredibly rare conformation that, when docked to, will give a higher score than anything in a smaller dataset. Without correctly accounting for such a conformation’s rarity, this will trend toward worse results with increasing sampling. Phrased differently, the best score approach is an outlier detector that is only in the correct ballpark when—by happenstance—no outlier conformations have been sampled yet. In contrast, more data from longer initial simulations should cause PopShift’s estimate to converge as the simulations do. The aforementioned outlier conformations, when correctly weighted by their rarity, will simply contribute to the overall picture of the ensemble instead of dominating the prediction.

To test this notion, we truncated our dataset as a series of fractions—that is, we took the first X% of each trajectory, where X corresponds to the fraction shown—and reran our analysis. Each result based on truncated data was generated by reworking that dataset as though it were the full length dataset, including featurization and clustering. We then inspected Pearson and Spearman correlations as a function of dataset size (Fig. 3). As before, we selected 20 frames per state to generate these truncated datasets. We also held cluster count fixed at 75, so that the total number of structures to dock to was not changing—only their diversity as the underlying feature trajectories became more mature and the quality of the MSMs providing the equilibrium population estimates.

**Figure 3.**
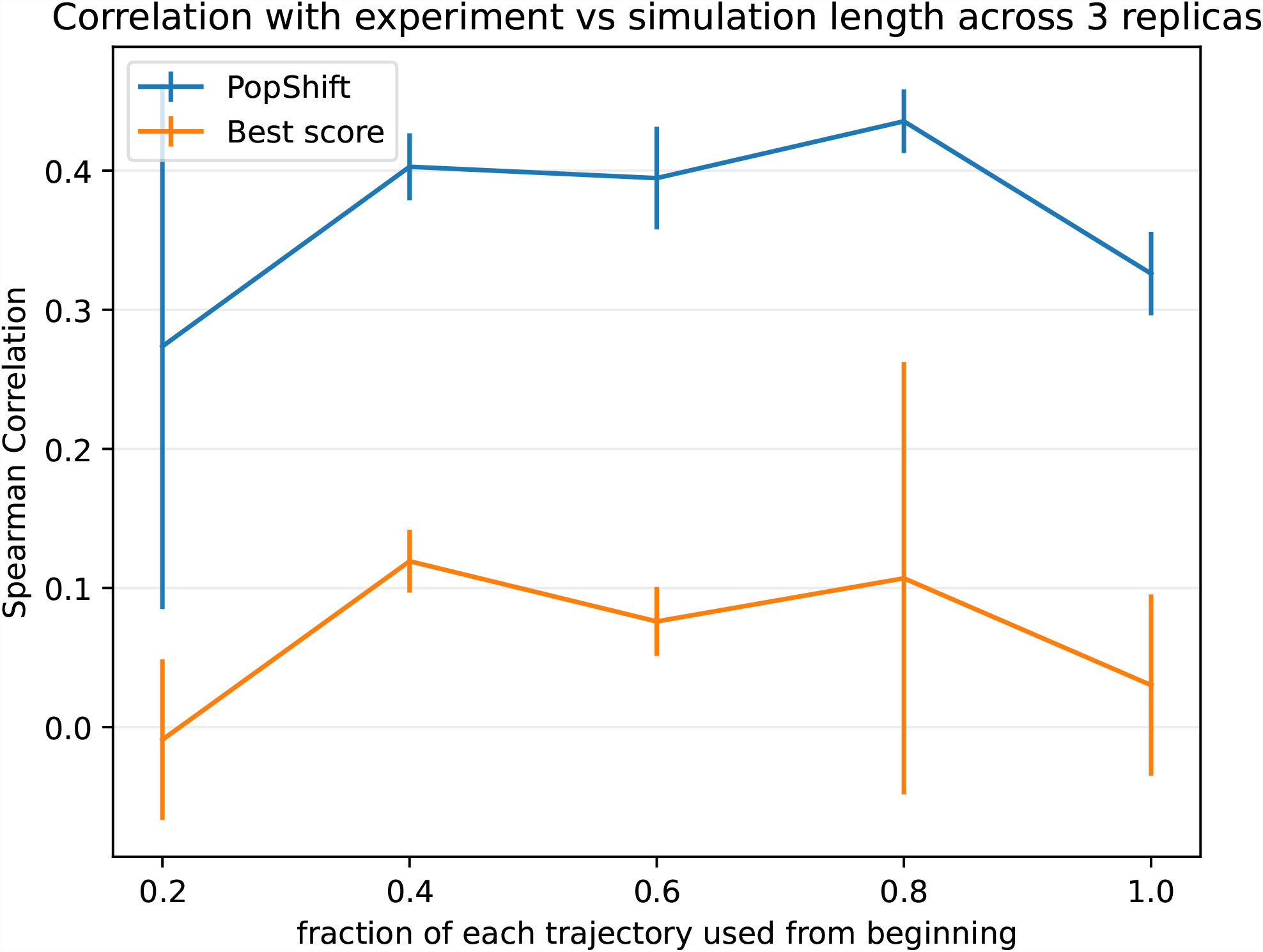
PopShift performs well as one varies the amount of simulation data, whereas traditional ‘best score’ ensemble docking gives worse performance and has greater statistical variation as data is added. The dataset used for MSM construction were truncated by taking the indicated fraction from the beginning of each trajectory. This can be viewed as asking the question, ‘what would happen if simulations had been stopped early?’ The error bars arise from the standard uncertainty in the mean across the three replica datasets.

We found that PopShift is less sensitive to the extent of input data than best-score ensemble docking. Because the number of structures docked to is constant across this sweep, it highlights how additional structural heterogeneity is not correctly indexed by simply looking for the most favorable score. As we suspected, best score gets worse with more data because it detects outliers. In other words, if an ensemble is scored by its most favorable possible interaction with ligand, the strain or unfavorability of that conformation on the protein is neglected. If one could create an ideal binding site for a ligand by moving residues out of the way such that it has ideal contacts, it would get very favorable docking scores when that conformaiton was docked to, but in fact the affinity for this site would be quite low because of how badly strained the protein would be by such rearrangements. In contrast, PopShift benefits from having more data, both in terms of the mean correlation with experiment and the statistical certainty in the results These results also emphasize that–at least for macroscopic binding constant estimation–our results for PopShift are not particularly sensitive to the length of input simulations. This is consistent with prior results suggesting that thermodynamic properties of MSMs converge quickly.^26^ Taken together, this implies that adding conformational heterogeneity to a docking campaign is best done by including a correctly weighted sample of receptor conformations, if the objective is ranked estimated affinity prediction.

### II.B PopShift retrodicts ligand poses, and their relative abundance

Given the low RMSE between PopShift’s predicted binding free energies and experimental measurements, we hypothesized that the approach also accurately predicts the pose the ligand adopts. specifically, we reasoned that any ligand likely adopts a wide variety of different poses in different protein conformations from the ligand-free MSM. If PopShift works as intended, protein-ligand structures where the ligand resembles ligand-bound crystal structures should have significant increases in their equilibrium probabilities compared to the same protein structure in the ligand-free ensemble. In this case, the distribution of RMSDs from the reweighted ensemble should be more favorable than the distribution from the original ensemble (i.e. using the state populations from the ligand-free MSM instead of updating the populations based on the strength of the interaction between protein and ligand)

To test our hypothesis, we compared the distribution of RMSDs to the ligand-bound crystal structure before and after reweighting the states based on the interaction with ligand (Fig. 4). For each sample from each MSM bin, i, we weighted its apo probability as being π_i_/n, where π_i_ is the equilibrium probability for that bin, and n is the number of samples drawn from each bin. We used Eq. 38 to estimate each sample’s probability in the presence of saturating ligand based on the docking score for that particular sample. We aligned based on pocket residue heavy atoms (residues within 5 angstroms of p-xylene in PDB 186L). RMSDs were then computed across all heavy atoms in the ligands. We plot both histograms for each replica in Fig. 4 to convey how reproducible the results are with different sets of simulations.

**Figure 4.**
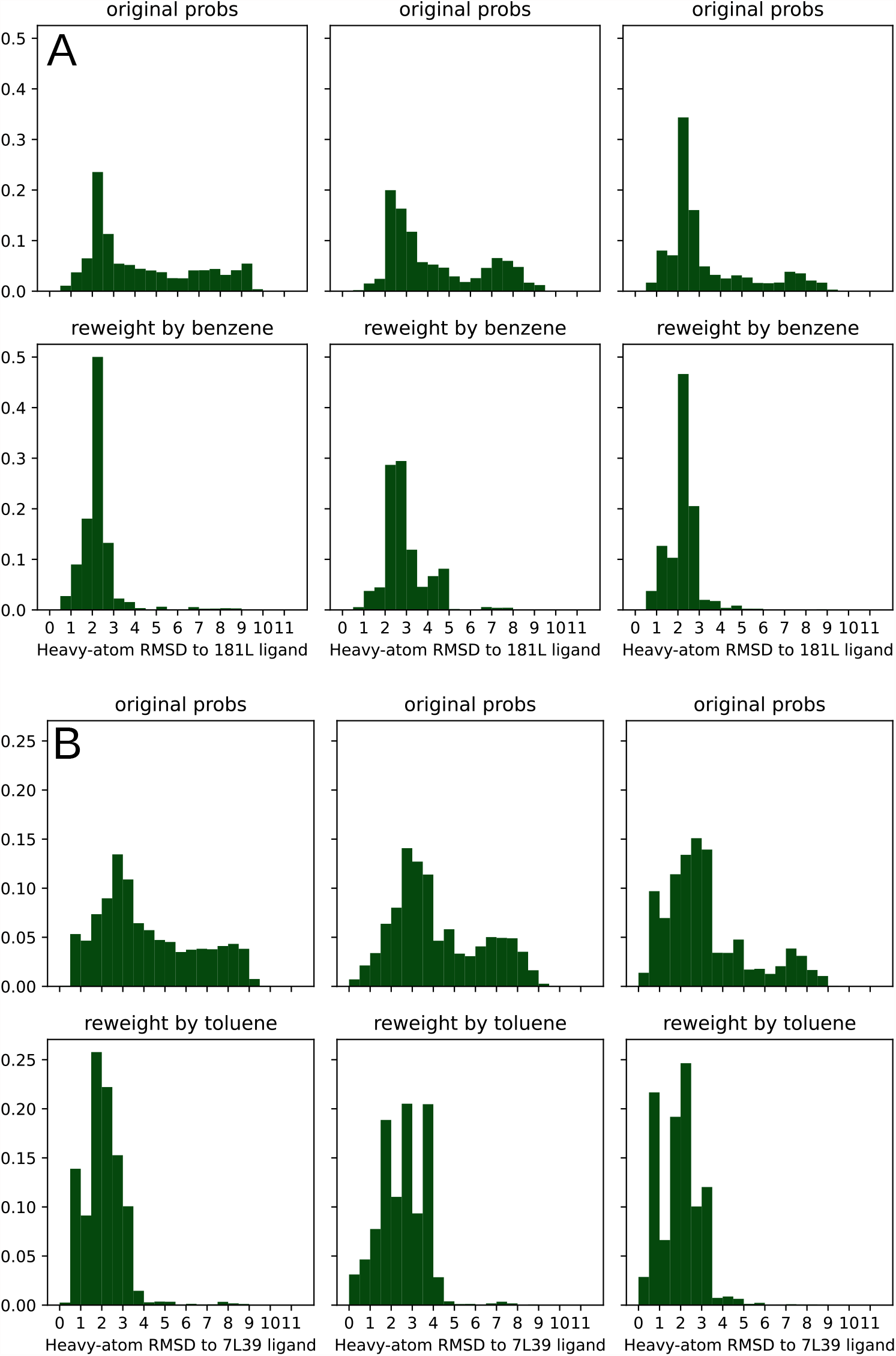
The population shift calculated by PopShift correctly favors ligand poses with a low RMSD to the crystal structure. The data histogrammed is the RMSD of the predicted pose to the holo crystal structure, where the structures are superimposed according to an alignment of their pocket atoms but not any ligand atoms. The RMSD histogrammed here is across all heavy atoms after this alignment. For each main panel, the three sub-panel columns represent individual replica datasets. The top row provide the ligand-free equilibrium probabilities and the lower row shows how the population is redistributed in the presence of ligand. Main panel A provides the data for benzene, while main panel B provides the data for toluene.

Our results for benzene and toluene show that PopShift does indeed favor low RMSD states compared to the broad heterogeneity in pose RMSDs from the original ensemble (Fig. 4, panels for ligand-free populations.) Many states from the ligand-free ensemble are not compatible with the experimentally observed binding pose, resulting in RMSDs between the best scoring pose and the ligand-bound crystal structure over 4 Å. When pose RMSDs were reweighted using pop-shifted equilibrium probabilities at saturating ligand concentrations, the distribution collapses and poses become holo-like. Interestingly, for some ligands such as toluene (Fig. 4 panel B), alternative conformations appear to be present. Given the way crystal structures solved at cryogenic temperatures are known to favor low energy structures and under-estimate structural heterogeneity, it is interesting to consider the possibility that the heterogeneity in poses that PopShift predicts for some ligands is real.

To test the generality of our results for the two ligands from Fig. 4, we devised a means to judge how closely our predictions agree with experiments across multiple compounds. Because experimental techniques have a hard time describing conformational heterogeneity, it is possible that poses dissimilar to experiment have relevance for the thermodynamic ensemble of the complex. Thus we chose to use three categories: one for configurations similar to the crystal pose, one for conformations that were dissimilar but likely still in the pocket, and one for conformations with clashes or completely alternative ligand placements. We reasoned this would be reasonably measured by aligning the receptor pockets, but transforming the ligands by that alignment.

We binned our histograms into three categories—fraction of samples that are similar to the crystal structure’s pose (RMSD < 2 Å), others that are in some alternative pose but probably still in the binding site (2 ⩽ RMSD < 4 Å), and ones that are likely in a very different pose or outside the binding site altogether (RMSD ⩾ 4 Å) in Fig. 5. We aligned the α-carbons of the pocket residues we used to build our MSMs, then transformed our predicted ligand poses by that alignment transform. Thus, high-RMSD scores likely emerge from poses that have significant displacements in center of mass—that is, poses that are not properly in the binding pocket. We named the three categories of poses ‘crystal-like’, ‘alternative’, and ‘miss’, as abbreviations of this interpretation.

**Figure 5.**
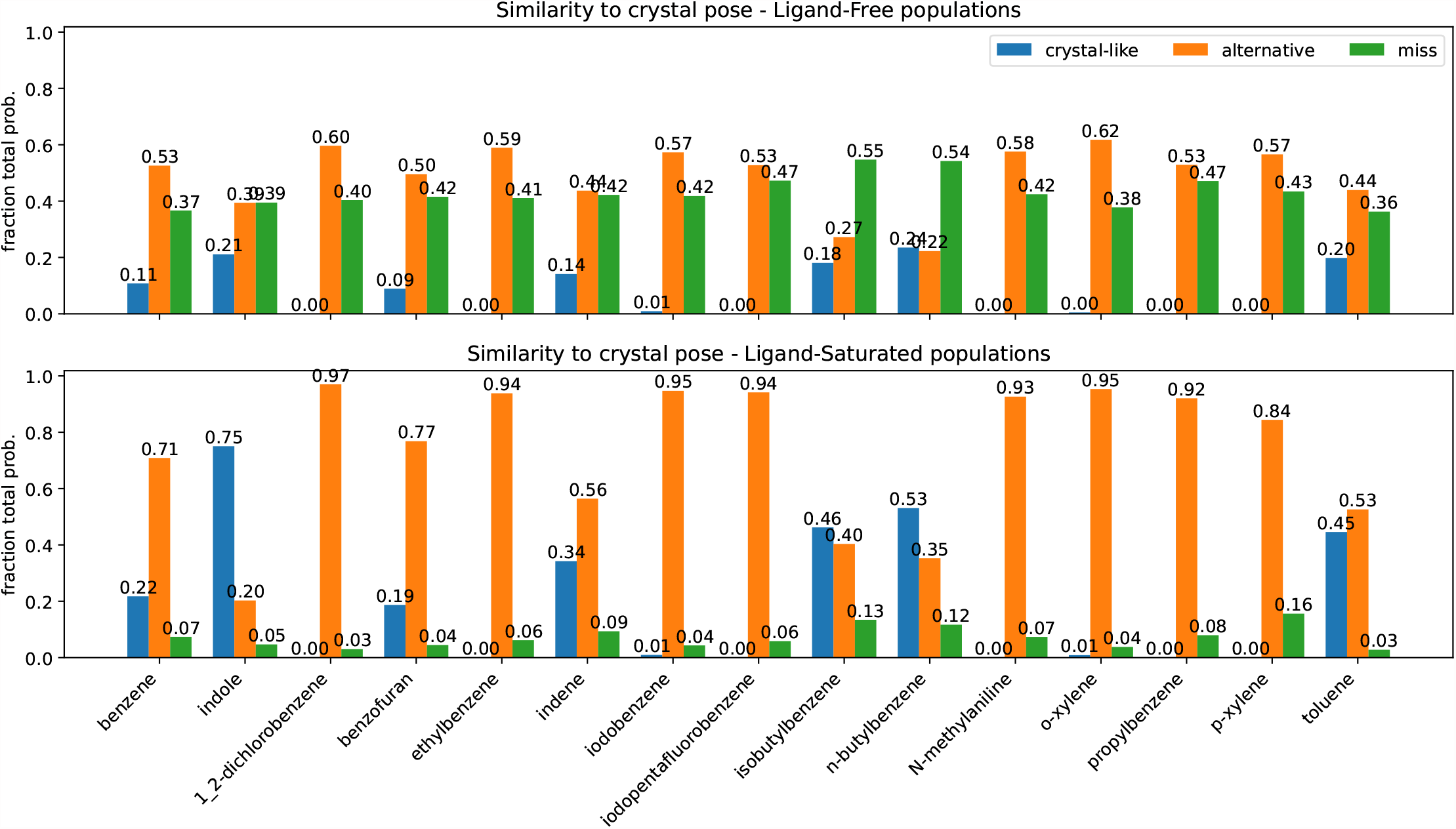
Summary of pose accuracy across all ligands studied. Each grouped bar represents the fraction of ligand poses that fall into the categories ‘crystal-like’, for within 2 angstroms of holo crystal structure, ‘alternative’, for between two and four angstroms RMSD from the holo crystal, and ‘miss’, for poses above four. As before the top panel represents the poses with an ‘apo’ ensemble weighting. The lower panel provides pop-shifted reweights for the same dataset.

The pattern we demonstrated for benzene and toluene in Fig. 4 is consistent across all the ligands we tested (Fig. 5). Poses we categorize as ‘miss’ are quite common with apo weights, but become rare after reweighting with PopShift. With ligand saturated weights, we often observe alternative poses. It is hard to know if these conformations exist in solution, but they are probable in the ligand biased ensemble, suggesting that they contribute nontrivially to our estimates of affinity.

### II.C PopShift predicts how ligands change the abundance of protein conformations

Because macroscopic affinities estimated with correctly weighted per-state affinities seem accurate, we reasoned that reweighted state probabilities might also be usefully accurate. We knew that Valine 111’s dihedral angle is able to occupy several rotameric states in apo simulations, but that the distribution is different upon ligand binding.^25,27^ Thus we hypothesized that the broad distribution from our ligand-free MSM should collapse to the binding-compatible one upon reweighting with ligand-saturated populations.

We tested this hypothesis by histogramming valine 111 angles from the receptor structures we sampled, weighted by both apo and ligand saturated state probabilities (Fig. 6). As in Fig. 4, we plot both histograms for each replica to convey the reproducibility of our results

**Figure 6.**
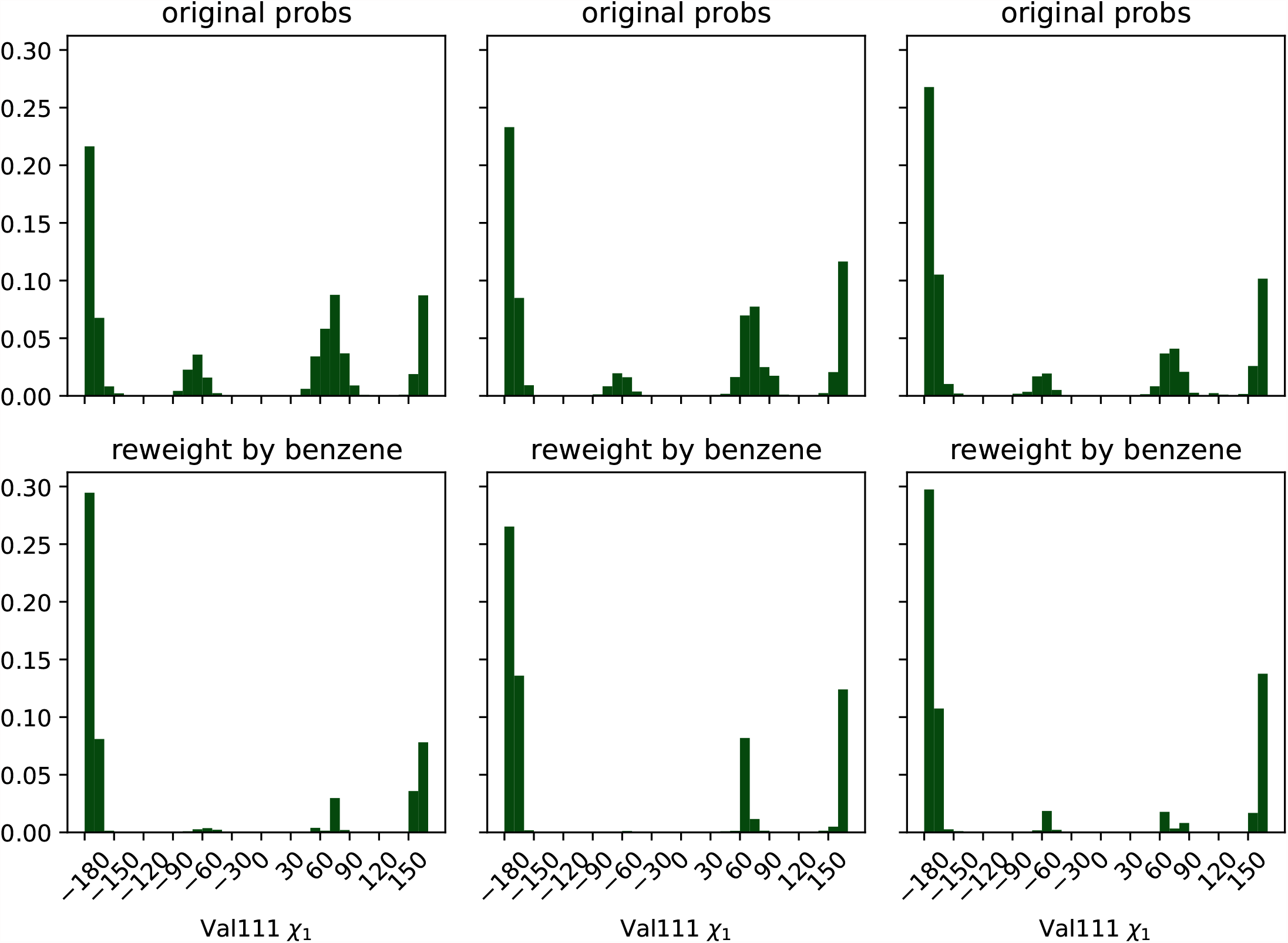
Three alternative conformations of the Val111 Chi1 angle in the ligand-free MSM collapse to mostly trans population in the presence of ligand, in agreement with the dominant pose seen crystallographically. The top row of plots are histograms of the Chi1 torsion across the frames sampled from the MSM states, weighted using the ligand-free MSM equilibrium probabilities. The second row displays the same data, but weighted by the benzene-saturated equilibrium probabilities. Each column represents the results from one fully independent replica.

We find that the ligand-free protein broadly populate several different structures, but the ligand shifts the population to favor the trans state, with some population of gauche +. This angle is noted as having many different distributions in RT crystal structures for liganded T4 lysozyme L99A protein.^25^ Trans is the angle modeled into cryo-structures from previous efforts (PDB ID 181L, 4W52).^22,28^ Our plots suggest that, like room-temperature X-ray structures, the ligand-saturated ensemble is heterogeneous, but does prefer certain angles, the primary of these being shared with the cryo X-ray structures. Thus the receptor population has shifted through conformational selection to a binding-compatible ensemble.

### II.D PopShift can estimate K_D_ and other ensemble features as a function of ligand concentration

Because the histograms from Fig. 4 represent unliganded and saturated conformational preferences, respectively, we hypothesized that inspecting conformational preference as a function of ligand concentration might help with analyzing binding preferences and allosteric effects. We wanted to know at what ligand concentrations certain histogram populations become more prominent since structural features not directly corresponding to ligand binding are often relevant for drug development—particularly in the case of allosteric modulators. For example, we previously identified both activators and inhibitors that bind a cryptic pocket in the protein TEM β-lactamase Looking at what structures are stabilized/destabilized by a ligand could provide a facile means to predict their effects on the structural preferences of distant sites and, ultimately, on function If a structural feature were used as a heuristic for some mechanistic action, that feature could be used to compute an EC50.

To display this transition, we computed ligand-rmsd histograms at various ligand concentrations and stacked them by descending ligand concentration, as a dilution series. To do this, we re-computed the histograms from Fig. 4 using Eq. 38 with a range of concentrations plugged in for x. Each histogram-bin’s probability was displayed using color, so that each row in the heatmap corresponds to a particular RMSD histogram at a particular ligand concentration. Bins, and therefore the X axis, match those from Fig. 4. We covered concentrations ranging from nM—essentially no ligand for these relatively weak binders—to 0.1 M—completely saturating for all the ligands we tried. Our predicted ligand dissociation constants are marked by an orange line.

We found that the populations of conformers outside the binding site were abundant until within an order of magnitude or so of the molecule’s K_D_, where a shift happened to structures that more closely resemble the ligand-bound structure. The poses closer to the K_D_ on the low concentration side contain a mixture of bound-like and ligand-free-like structures. They are similar to the ligand-free-weighted distribution we see in Fig. 4.

## III Conclusion

In this work we have presented PopShift, a framework for estimating binding free energies in a manner that correctly weights the conformational heterogeneity present in the ligand binding sites of proteins. PopShift’s estimated binding free energies from docking scores perform well compared to other common methods for the simple problem of lysozyme L99A binding to small organic compounds. Our results demonstrate that adding receptor fluctuations into docking is indeed a significant improvement over making the simplifying assumption that a single protein structure encodes all the relevant information. Further, our approach provides an approximation of the receptor-ligand complex ensemble, which has utility in mechanistic studies, such as those focused on tuning the abundance of receptor conformations known to correlate with function.

The future directions for this approach are many. PopShift of docking scores for affinity estimation is still limited by the performance of the docking scoring function. Thus it is likely that applying PopShift to more challenging problems, such as for ligands with many rotatable bonds or charge, will require more sophisticated estimates of per-state K_D_s. Using per-state estimates from Generalized-Born or Poisson Boltzmann rescoring, or absolute or relative binding free energy simulations, is therefore an exciting and immediate future direction for this work.^29^ More broadly, we see opportunities to apply this framework to other perturbations to an MSM sampled from one thermodynamic state, without having to redo the sampling in those new states—such as gracefully integrating multiple protonation states for either ligand or receptor, and indexing the relative free energy changes of mutations. Expansive and expensive sampling, done once for some reference model, can thus be reweighted to solve a host of important problems facing modern Biophysics.

## IV Methods

### IV.A PopShift formalism

We noted that we could treat an MSM as a system’s partition function, since the equilibria between each state and the equilibrium probability, or population, of each state is available. Thus, we supposed that the binding polynomial formalism of Wyman and Gill [18] would give us traction for describing how binding affects this representation of the system’s ensemble. The discrete treatment of state space seemed natural in the context of an MSM-representation of an ensemble. For the sake of completeness, an equivalent derivation based on the Zwanzig perturbation formula is included in Appendix A.

Assume we have an MSM representing an ensemble discretized into n states where the abundance of the *i*^th^ state is π_*i*_, its equilibrium population. Suppose further that we can obtain an estimate of the affinity of a ligand or an array of ligands to each state, k_*ij*_, indexed by the i states and up to t ligands bound, the free concentration of which is given by χ_*j*_. Then we can write the partition function, Z, in terms of all such equilibria available to the system as in Wyman and Gill [18]:

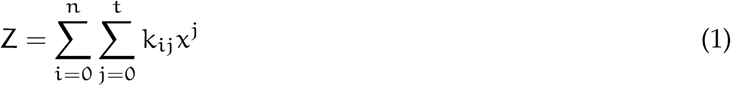

Starting from here has several nice features; it is clear how we might extend this formalism for multiple ligand binding sites, or for multiple ligand species by adding a third index to Eq. 1. We can rearrange this definition to obtain an expression for the fractional population of the various allosteric states. We’ll express this in terms of the sub-binding polynomial, P_*i*_.

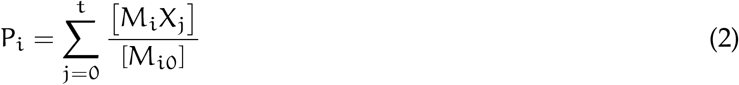

Here, [M_*i*_X_*j*_] is the macromolecule-ligand complex in state i with j ligands bound, and [M_*i*0_] is the concentration of that state in the absence of ligand. We’re considering equilibria relative to the abundance of the allosteric conformation sans ligand. We’ll abbreviate the equilibrium constant between each ligand free state and the reference state as: 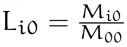. Thus P_i_ becomes:

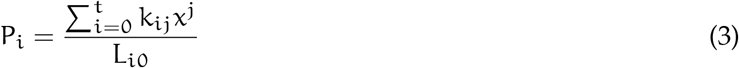

And then:

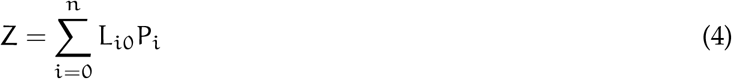

If we would like to consider the reference condition to be all unligated species, we can define a normalized binding polynomial, P, using this expression for Z:

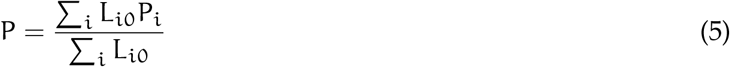

Since the fraction of the *i*^th^ form, α_*i*0_, can be written in terms of un-ligated equilibrium constants:

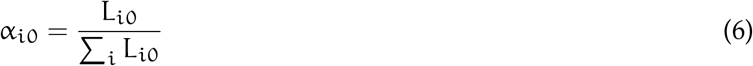

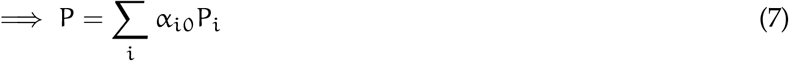

Noting that these α_*i*0_ are exactly the equilibrium probabilities from our unligated MSM, let *α*_*i*0_ → π_*i*_. Finally, we can write an expression for the population of the *i*^th^ state under the influence of ligand, 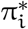, as:

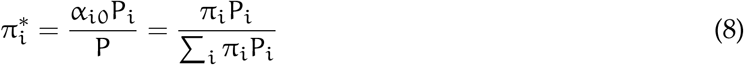

### IV.B Calculating ligand binding free energies for one monovalent site

Starting from the definition of a one ligand binding equilibrium constant, we can obtain an expression for the free energy change associated with that binding constant in terms of the binding polynomial corresponding to this. Writing the macro equilibrium constant, K, of all macromolecule-ligand complexes with one ligand bound as [MX] with [M] and *x* as the free receptor concentration and ligand activity, respectively:

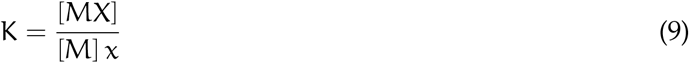

Note that this is an association constant; its reciprocal would be the dissociation constant. Thus the degree of binding, 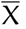, is:

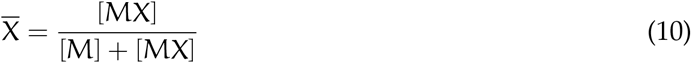

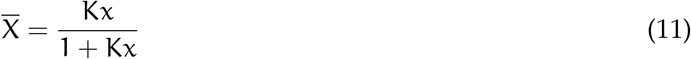

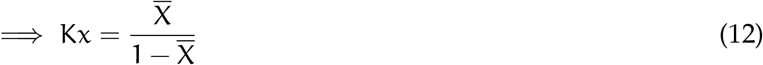

From here we can get to the binding free energy, which is by definition the ligand concentration at which the fully ligated and unligated species are equal. In the binding polynomial literature this is known as the ‘median activity’, *x*_m_.

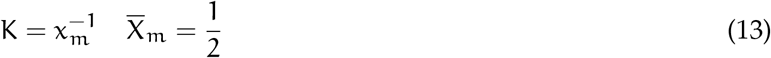

We can express the fraction bound as the derivative of the binding polynomial. For our simple one site one affinity model, the binding polynomial becomes:

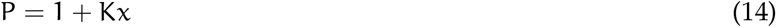

The binding curve, 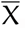, can be recast as the derivative of the log of P with respect to the log of x:

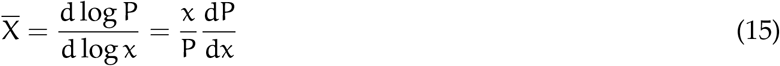

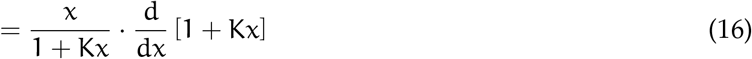

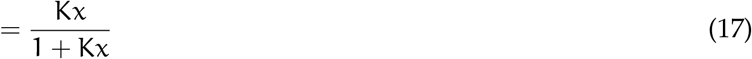

This expression is general; for any binding polynomial describing a system the binding curve can be obtained by writing the derivative of the binding polynomial with respect to the log of the ligand activity. If there are multiple ligands, then the fractions bound can be described in this way using partial derivatives with respect to the log of the various ligand activities.

From here we can also consider a free energy of binding, with an eye toward a more general expression (where multiple ligands might bind). Since we have derivatives of the binding polynomial, we should also be able to write integrals of these to get to the expressions we started with.

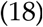

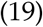

Where *A*_*u*_ is the area under the binding curve.

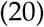

Also note that, as mentioned before, P^−1^ = *α*_0_, is the fraction of unligated macromolecule.

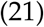

Likewise if we integrate from maximum ligand activity to a particular log ligand activity, we can write the area above the curve. This will be the fraction of fully ligated macromolecule (the species with all t receptor binding sites occupied).

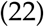

*α*_*W*_*efindthatPopShiftismoreaccuratethancommonstrategies,suchasdockingtoasinglestructureandtraditionalensembledocking*

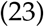

The overall binding constant is determined by this value, since it is the ratio of the unligated species to the fully ligated species.

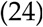

Picking the median activity here means the above expression reduces to:

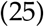

For one site, t = 1.

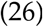

The standard free energy change for this system is the work required to add an infinitesimal amount of ligand from a reservoir at standard state to the macromolecule. This can be expressed in terms of chemical potentials as a variation of the free energy:

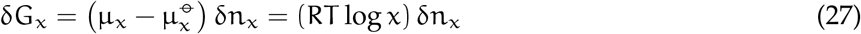

Where G_x_ is the free energy of ligand binding, δn_x_ is an infinitesimal amount of X, μ_x_ is the chemical potential of X, and 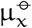 is the standard state chemical potential of X. We can consider how much work it would take to saturate a mole of our receptor by integrating this formula across our degree of binding formula, and asserting that 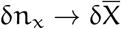.

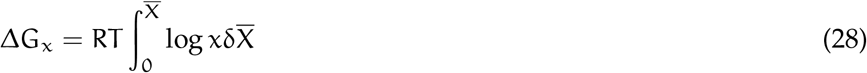

Since this work is the work of going from M + tX → MX_t_, we can see the substitution for the median activity that gets us to the customary relationship.

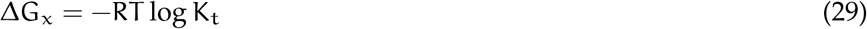

#### IV.B.1 One binding site, multiple conformations

Now we’re positioned to ask what happens if we have multiple conformations that each have a particular affinity for a ligand, assuming that the affinity is arising from one binding site. The formula for the multiple-conformers binding polynomial is:

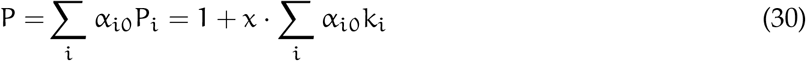

Note that we’ve already started from a model where we’ve assumed many states with one binding site for one ligand, with each state having its own binding constant. Then the Adair constant for the ligated state, K_1_, is the macro-equilibrium constant for the reaction.

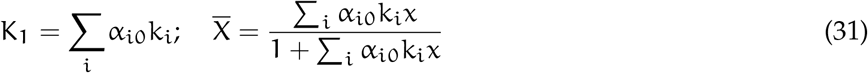

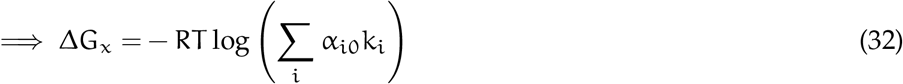

Using the *conformational selection assertion* to set the state fractions to the equilibrium probabilities for each MSM bin from our model, *α*_*i*0_ → π_*i*_, and each of the 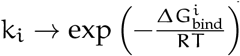. These can be had by docking, alchemy or other means that produce binding constants or free energies.

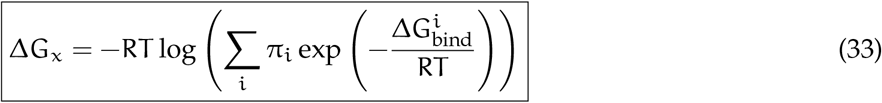

### IV.C Population shift per MSM bin

Based on the expression for the reweighted state probabilities above, Eq. 8, we would like to write an explicit formula for our single site, multiple conformation case. To do so we write the sub-binding polynomial as before, with k_*ij*_ the equilibrium between the *i*^th^ state with j ligands bound and the reference state M_*i*0_:

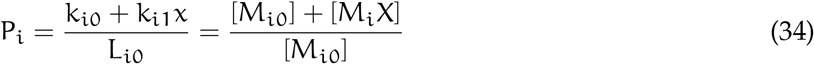

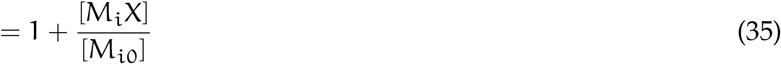

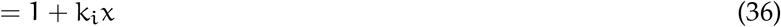

From here we substitute back into Eq. 8, the expression for 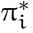:

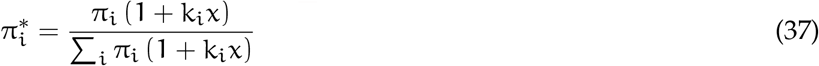

Let 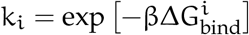, the micro-equilibrium constant estimated for that state.

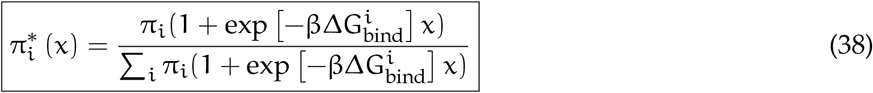

Note that Eq. 38, the PopShift equation for a single site binding model, has the correct limiting behavior. At high x, the populations of the states are dominated by the favorability of binding to each state because the right hand numerator term and the denominator both grow with x, but the left-hand term does not. Conversely, at very low *x* the state probabilities are very nearly the apo ones, as the right hand term in both the numerator and denominator becomes small relative to the left-hand term.

### IV.D Lysozyme MSM simulations and construction

#### IV.D.1 Simulations

Simulations were run with Gromacs.^30^ Three sets of ten simulations were run starting from protein coordinates taken from PDB ID 187L using the Amber03 force field.^31^ Five of the trajectories in each set totaled 4μs of sampling, while the other five totaled 8μs. The protein was solvated with TIP3P explicit water in a dodecahedral box that extended one nm beyond the protein in any dimension and eight chloride ions were added to neutralize the charge.^32,33^ This system was energy minimized with the steepest descent algorithm until the maximum force fell below 10 kJ mol 1 min 1 using a step size of 0.01 nm and a cut-off distance of 1.2 nm for the neighbor list, Coulomb interactions and van der Waals interactions.

The system was then equilibrated at 298K in a 1 ns NVT simulation followed by 1ns NPT simulation with a position restraint on all protein heavy atoms (spring constant 1,000 kJ mol^-1^ nm^-2^). A long-range dispersion correction was employed for both energy and pressure. All bonds were constrained with the LINCS algorithm.^34^ Cut-offs of 1.2, 0.9 and 0.9 nm were used for the neighbor list, Coulomb interactions, and Van der Waals interactions, respectively. The Verlet cut-off scheme was used for the neighbour list and particle mesh Ewald was employed for the electrostatics (with a grid spacing of 0.12 nm, PME order 4, and tolerance of 1e−6.^35^ The v-rescale thermostat (with a time constant of 0.1 ps) was used to hold the temperature at 298 K and the Berendsen barostat was used to bring the system to 1 bar pressure.^36,37^ For the production runs, the position restraint was removed and the Parrinello-Rahman barostat was employed.^38^ Snapshots were stored every 10 ps. Structures were visualized with PyMOL, and trajectories with both PyMOL and VMD.^39,40^

#### IV.D.2 MSM construction

MSMs were constructed using Deeptime independently for each set of 10 simulations.^41^ Clustering data was managed using the `RaggedArray` class from `enspara`.^42^ Backbone and all 𝒳 dihedrals for any residues with heavy atoms within 5Å of p-xylene in PDB 187L were selected as input features. This feature space was reduced using TICA,^43^ with lag times of 1, 2, and 5 ns and with a kinetic variance cutoff of 0.9 using commute mapping. We used k-means to cluster this reduced feature set, chosing our number of states using the cross-validation approach taken by Meller et al. [23]. Briefly, the reduced features were clustered by splitting features into 10 train-test pairs, where k-means with a range of k was used to cluster only the training set. Test set trajectories were then assigned to clusters using euclidean distance to the k centroids resulting from the ‘training’ clustering. MSMs were fit to the train and test pairs using the MLE method.^44^ The first 10 eigenmodes of both models were then VAMP-2 scored^45^ using the train model, to estimate how over-fit the model was to cluster count.^**pande2015**, 46^ The number of clusters chosen for final model fitting was the point at which the VAMP-2 score of the test data starts to decline, which was k = 75 in this case. The complete set of input features was reclustered using k-means with 75 clusters and then an MLE model was fit with a lag time of 20 ns, after scrutinizing the implied timescales of the data with various lag times.

### IV.E PopShift workflow

The workflow used here to do the PopShift post-processing of our ensemble docking run is as follows :

1. Obtain a satisfactory ligand-free MSM.^a^
2. Sample a number of receptor conformations from each bin of the MSM using the assignment trajectories from clustering (Frame-picking).
3. Align these conformations so that they will fit neatly in a docking box.
4. Dock to each sample, saving the ligand pose and docking score.
5. Compile docking scores into free energies of binding using Eq. 33 and the equilibrium probabilities from the ligand-free MSM.
6. Compute reweighted state populations from the docking scores using Eq. 38.

#### IV.E.1 PopShift implementation

To pick frames, the assignment trajectories used for model selection were sorted into lists of frames corresponding to each cluster center, and then several of these (20 for the data in Figs. 2 and 3) were selected in uniformly random fashion. These frames were extracted from the coordinate trajectories and iteratively aligned^47^ by the *α*-carbons of their pocket residues (defined in the same fashion as for MSM construction in Sec. IV.D.2), using LOOS.^48,49^ Ligands and receptors were prepared using `prepare ligand.py` and `prepare receptor.py` from AutoDock tools.^50^ We parallelized the preparation process by using GNU `paralle`l.^51^ Docking was performed using a box with 12-angstrom sides centered on the centroid of the average structure of the aligned frames using SMINA.^24^ Each docking run targeting each extracted conformation was performed as an independent single CPU task using Jug.^52^ We used the SMINA and Jug versions hosted on conda forge. For SMINA the binary we used returned the version statement (from calling `smina --version) Smina Nov 9 2017`. `vBased on AutoDock Vina 1.1.2.` For Jug the version statement returned by the python module was `2.2.2.` No modifications were made to the docking energy model for this study. Docking scores were extracted and collated into ligand-indexed JSON associative arrays using scripts provided in the PopShift package. PopShift is available as open-source software and can be found on the Bowman Lab Github: https://github.com/bowman-lab/PopShift

### IV.F Disappearing ligand absolute binding free energy simulations

#### IV.F.1 Starting pose selection

The binding modes of the ligand for free energy calculations were selected using 5 methods. For ligands with a known crystal structure bound to T4-Lysozyme, we selected the MSM pose with an RMSD closest to that of the crystal structure. All poses had an RMSD < 2Å to the crystal pose and thus would be considered the same binding mode.^54^ The exception to this was 1,2-dichlorobenzene which did not have an MSM pose that closely matched the known crystal structure. In the case of 1,2-dichlorobenzene, we used the exact pose from the crystal structure.

For ligands without a crystal structure, we took a known crystal structure most similar to the ligand and posed our ligand accordingly in several poses. Each pose for each ligand was simulated for 2 ns and an RMSD analysis of the ligand throughout the trajectory as compared to the starting pose was run. The pose with the smallest change in RMSD and the smallest variance in RMSD was chosen as the most stable pose, and was the pose used for the remainder of calculations. We overlaid 2-ethyltoluene with the crystal pose of o-xylene, and then flipped 2-ethyltoluene so the ethyl group and methyl group would align first with the 1-methyl and 2-methyl of o-xylene as pose 1, and the 2-methyl and 1-methyl as pose 2. We overlaid 3-ethyltoluene with the crystal pose of o-xylene. We aligned the ethyl group of 3-ethyltoluene with each methyl group of o-xylene resulting in 4 different poses. We also aligned the methyl group of 3-ethyltoluene with each methyl group of o-xylene resulting in 4 more different poses. We overlaid 4-ethyltoluene with the crystal pose of p-xylene, and then flipped 4-ethyltoluene so the ethyl group and methyl group would align first with the 1-methyl and 4-methyl of o-xylene as pose 1, and the 4-methyl and 1-methyl as pose 2. We overlaid thianaphthene with crystal pose of indene as pose 1, and flipped the thianaphthene across its length so the sulfur would be on the opposite side as pose 2.

Lastly, for m-xylene, we began with a process similar to that of 3-ethyltoluene. We overlaid m-xylene with the crystal pose of o-xylene. We aligned the methyl groups of m-xylene with each methyl group of o-xylene resulting in 4 different poses. Again, an RMSD was used to choose the most stable pose to use for the remainder of calculations. However, upon free running free energy calculations, we observed m-xylene switch to a different stable pose after 1.5 ns, impacting the free energy calculation. For m-xylene, the stable pose found at 1.5 ns into the free energy calculation was chosen.

#### IV.F.2 Ligand and protein parameterization

The ligands were parameterized with Open Force Field version 2.0.0 and charged with AM1-BCC charges.^**mobley2022**^ The protein (PDB 7l38) was prepared using OpenEye Spruce to add hydrogen atoms at pH 7.0. The protonated protein was then parameterized using AMBER ff14SB and the TIP3P water model was used for the waters. GROMACS was used to solvate and add a salt concentration of 150 mM to the ligand and protein-ligand systems. Each ligand system was energy minimized and NVT equilibrated, then a 2 ns NPT production run was performed. Each protein-ligand system was energy minimized and NVT equilibrated, then a 2 ns NPT production run was performed. The trajectory of the production run was used to select the atoms and dihedrals for the Boresch restraints to restrain the ligand to the binding site during simulation.^53^

#### IV.F.3 Running Absolute Binding Free Energy calculations in GROMACS

Simulations were run using GROMACS 2021.2. For binding site simulations, we used 20 lambda windows. In this protocol, we first restrained the ligand to the binding site, turned off the coulomb interactions, then turned off the vdW interactions. For unbound ligand simulations, we performed absolute hydration free energies. In this protocol, we first turned off the coulomb interactions, then turned off the vdW interactions.

Prior to running production simulations, every lambda window was energy minimized for 5000 steps using steepest descent and equilibrated at constant volume for 10 ps at 298.15K. Production simulations were run for 15 ns per lambda window with an NPT ensemble. During production, replica exchange was attempted every 200 steps. See Fig. 9 for a graphical representation of this schedule.

**Figure 7.**
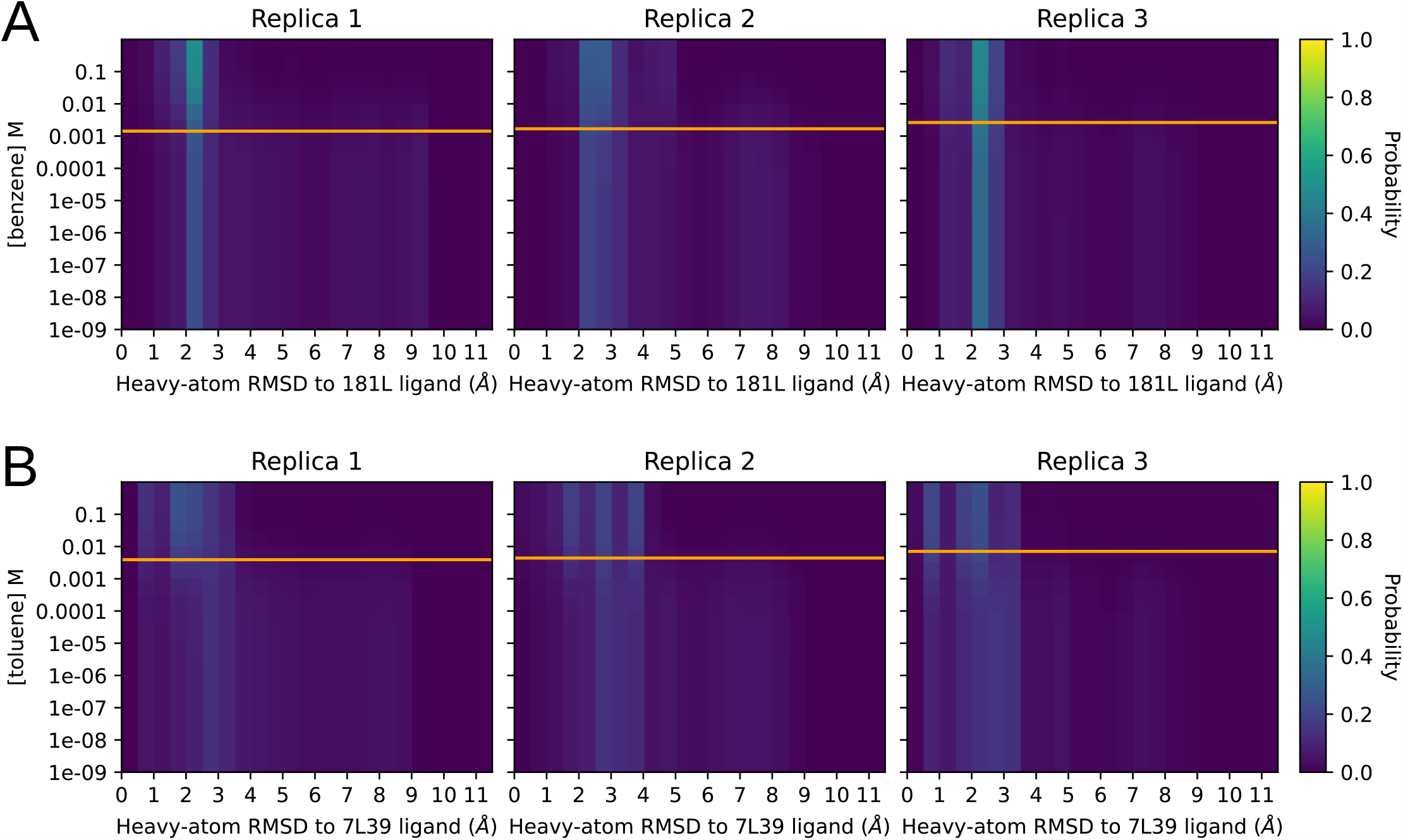
The distribution of RMSDs to a ligand-bound reference structure as the concentration of the ligand is varied. The Y axis is ligand concentration, and the X axis is the pocket-aligned Ligand RMSD to holo crystal structure, histogrammed as before (Fig. 4). Each row in each heatmap is from the same raw data being histogrammed with different weights, computed by plugging the Y-value matching that row into Eq. 38. The orange line on each replica plot shows the K_D_ as calculated for that replica using 33, and is shown for reference. The top row in each heatmap is holo-like, because it is at high ligand concentration, and the bottom row is apo-like because it is at low ligand concentration. Being able to titrate observables measurable from conformations in this way could provide exciting opportunities to understand ligand efficacy in systems with allosteric behavior.

**Figure 8.**
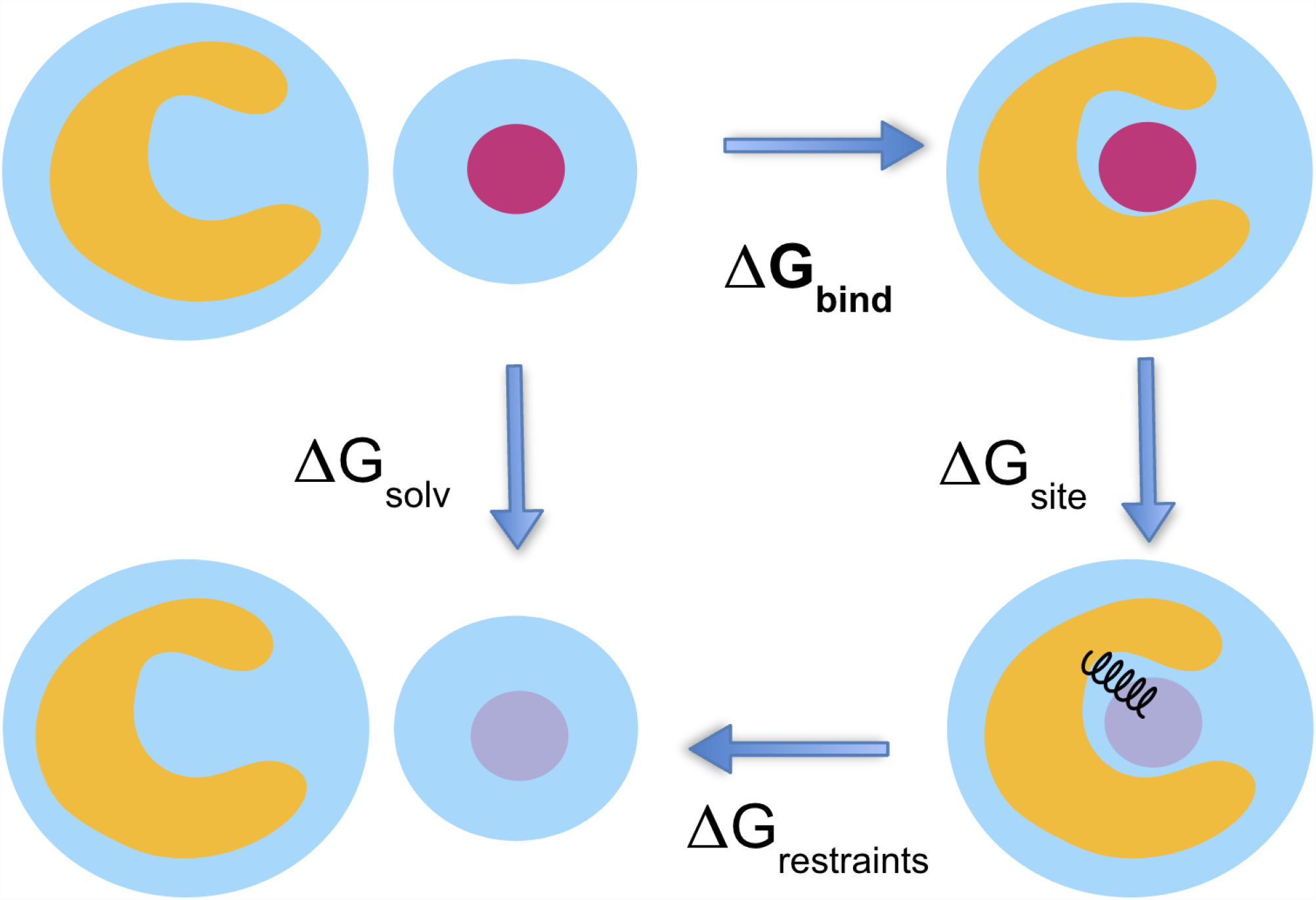
This cartoon depicts the thermodynamic cycle and transformation used for the absolute binding free energy calculations from Fig. 2. We are taking ΔG_bind_ = ΔG_solv_ − ΔG_site_ − ΔG_restraints_. In its weakly and non-interacting state, the ligand is free to leave the binding site. We use orientational Boresch-style restraints to reduce the phase space that must be sampled.^53^

**Figure 9.**
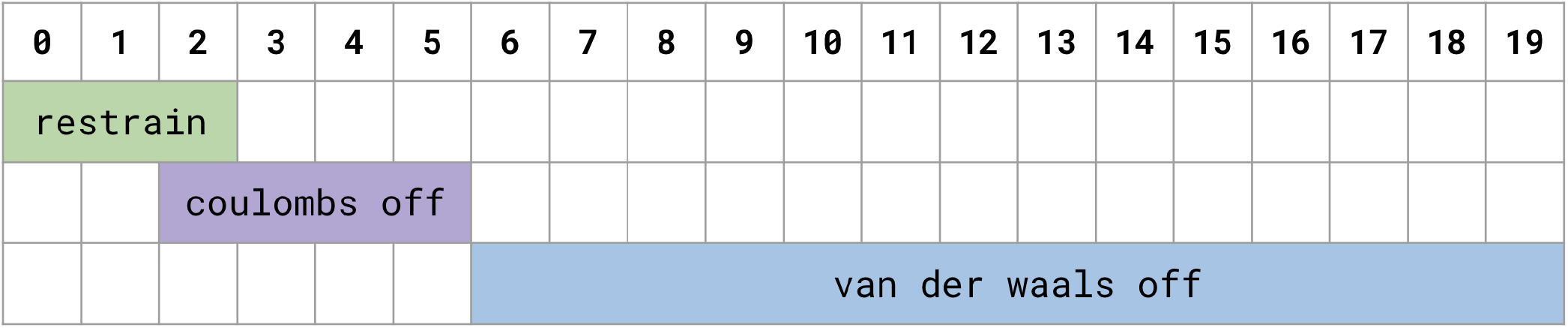
This λ schedule depicts the schedule for restraining and scaling down ligand interactions (coulomb and vdW) for the protein-ligand protocol discussed in the body text.

#### IV.F.4 Analysis of absolute binding simulation results

We obtained the free energy difference using the alchemlyb/pymbar package MBAR estimator. The first nanosecond of the 15 ns of production simulations were discarded as equilibration. Each ligand was inspected for symmetry and the trajectory of the protein-ligand system in its unrestrained state was inspected to determine whether all symmetries were equally sampled.^55^ The free energy difference of ligands with at least one axis of symmetry without adequate sampling of symmetries were corrected using the equation:

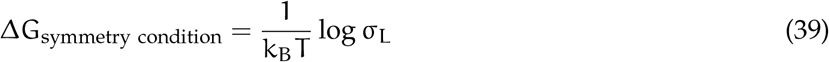

Where k_B_ is the Boltzmann constant, T is the temperature, and σ_L_ is the ligand symmetry.

## Software

Python command line tools written to perform the PopShift calculations discussed here are distributed as free software at: https://github.com/bowman-lab/PopShift

## Data

Data files and scripts used to analyze results and produce figures, as well as trajectory data, will be available upon request. The data will also be hosted (possibly via links or other download means, depending on which data it is) at https://github.com/bowman-lab/popshift-ms-data

## Acknowledgements

We thank Alan Grossfield for galvanizing discussions and encouragement. We also thank Artur Meller and Matt Cruz for insightful comments over the course of the work. We thank Justin J. Miller for helpful discussion of ITC reference data. This work was funded in part by NSF MCB-2218156, NIH NIA RF1AG067194, and the Basser Center for BRCA at the University of Pennsylvania, to G.R.B., and NIH R35GM148236 and R01GM132386 to D.L.M. G.R.B. holds a Packard Fellowship for Science and Engineering from The David & Lucile Packard Foundation.

## A Derivation of MSM-docking using the Zwanzig relationship

The binding of a ligand to a protein is described by the reaction

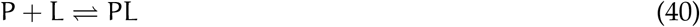

The binding free energy is related to K_D_, the dissociation constant, by:

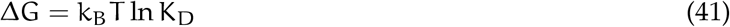

or:

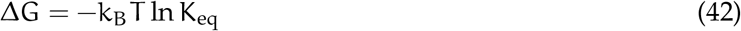

The binding free energy is computed from the ratios of partition functions describing the elements of the complex.

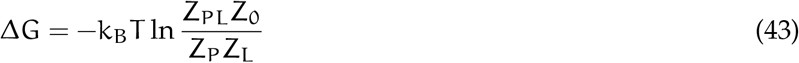

Here the various sub-partion functions are defined with Z, with the subscript indicating what they encompass. PL for the protein-ligand partition function, 0 for the solvent/milieu, and P and L for the protein and ligand separately. By splitting up the natural logarithm we can focus on hydration free energy and protein ligand binding of the expression separately:

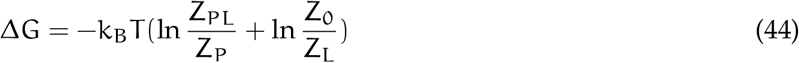

Where:

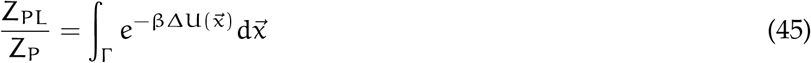

One of the limitations of binding free energy calculations is that the partition function integral goes over the full phase space Γ, the full conformational space accessible to the protein and to the ligand. In part to address this limitation, we can take the integral across the state space of either system by treating the transition from bound to unbound (state A to state B) as a perturbation.^19^ The perturbation formula is thus an approximation that becomes exact in the limit of infinite sampling.

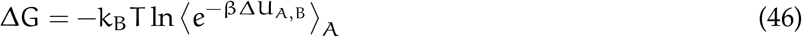

Where ΔU_A,B_ is the difference in energy between states A and B.

To combine exponential averaging with MSMs, we fully partition the state space, extract the equilibrium probabilities and calculate the differences in potentials between ligand free and ligand bound conformations.

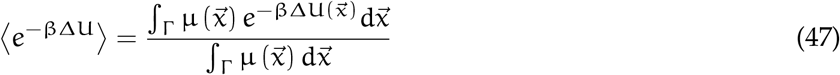

Where μ is the stationary distribution the transfer operator our msm approximates transports density toward, and 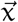 is a point in configuration space.^56^ Because of this discretization of configuration space, the integral from Eq. 46 can be split across the states of the system; if we discretize this integral across sub-volumes in configuration space this becomes

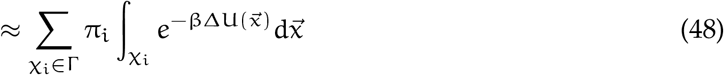

Here π_*i*_ is the equilibrium probability of the *i*^th^ state, and χ_*i*_ is the subset of Γ corresponding to that state. Taking these sub integrals per state could be done using alchemical free energy calculations restricted not to sample within each bin in phase space–that is, within each 𝒳_*i*_. Here we used binding probabilities estimated via docking to each state instead.

Suppose that we can obtain an estimated equilibrium constant for a given conformation using docking.

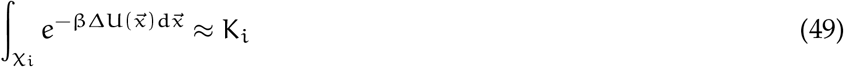

Where K_*i*_ is an equilibrium constant estimated from docking for the i^th^ state. Then we get the relatively straightfowrward:

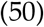

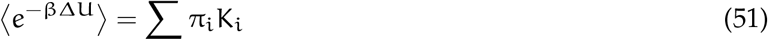

Plugging this expression back into Eq. 46 gives us:

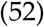

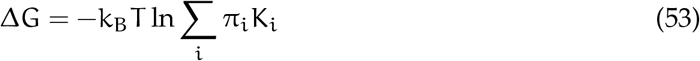

### A.A Using state populations to estimate EC_50_

A stronger or weaker overall binding of ligand may not be predictive of changes to macromolecular function. For example, it is possible for a ligand to bind to a known allosteric site quite tightly, but to have the mode of its binding fail to stabilize conformations that change its function.^57^ Put another way, it is likely, even) that all modulator-bound conformations don’t have the same potency; some such conformations likely matter more than others, and the affinity of the modulator to that conformation may not be the sole predictor of this activity. To avoid confusion with chemical activities, we will refer to this as *potence*.

Suppose that we define some measure of a state’s propensity to progress through some binding or reaction cycle called potence, 𝔓. This measure could be based on some criterion from the literature, experimental data, or some ansatz that, once formulated, could be checked to see if it recapitulates experimental data; for example states with feature Y are part of the (de)activated macrostate. If potence is a function of conformation, 𝔓 (M_*i*0_), then it can be calibrated relative to the unperturbed ensemble and the saturating one for the receptor’s unligated activity, and the equilibrium constants can be used to estimate an EC_50_ based on the modulated probabilities. In other words, even a rough heuristic that is only proportional to function may be sufficient to estimate an EC_50_. Starting with the unligated potence:

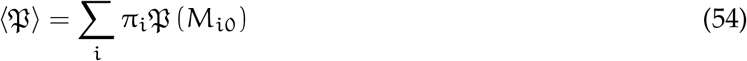

Using Eq. 38, we have a path to calculating the EC_50_.

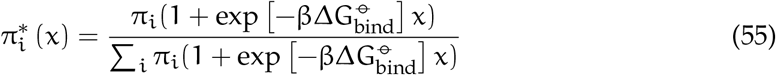

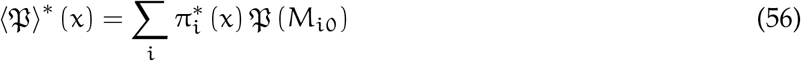

The EC_50_, should be the ratio of abundance of the potence score states equal to 1/2 its max value. Since we can simulate this by putting in a saturating concentration of ligand, we can write:

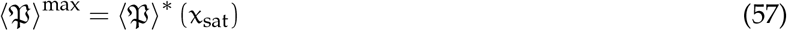

Then we can create a fractional potence score and set it equal to 1/2:

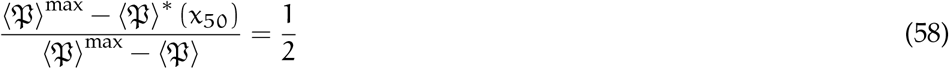

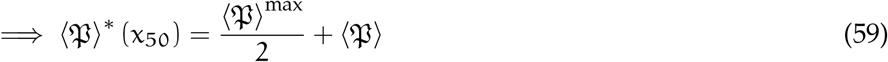

This expression can be solved numerically for χ.

## Footnotes

a Note that for thermodynamic observables all that is really needed for step 1 above is a discretized state space or clustering of input features, and an associated collection of equilibrium probabilities for each state. MSM construction, especially with a collection of shorter trajectories as we have, is a sensible path to obtaining this association, but others are possible.

## Notes

https://github.com/bowman-lab/PopShift

### Competing Interest Statement

The authors have declared no competing interest.

### Summary of Updates

Typos were caught, as was an issue with which dataset was being plotted in Figure 2. Several segments were rephrased after receiving more feedback from collaborators. Much of the rephrasing was done to increase clarity.

